# Limited Effects of Isolated Congenital Anosmia on Cerebral White Matter Morphology

**DOI:** 10.1101/2025.11.04.686499

**Authors:** Anja L. Winter, Moa Peter, Evelina Thunell, Johan N. Lundström, Fahimeh Darki

## Abstract

Lack of sensory input is associated with alterations in brain morphology; mainly in or near cerebral regions normally devoted to processing of the missing sense. We have in multiple studies demonstrated that the only consistent morphological finding within the gray matter of individuals born without the sense of smell (isolated congenital anosmia; ICA), are changes in or near the olfactory sulcus. For the connecting tissue of the brain, the white matter (WM), previous studies have yielded inconsistent findings. Here, we show that individuals with ICA (n=49) exhibit alterations in WM volume as compared to age- and sex-matched controls. Consistent evidence from both voxel-based morphometry and multi-voxel pattern analysis shows that individuals with ICA show decreased WM in areas surrounding the olfactory sulcus. Importantly, no WM alterations were found in areas surrounding the olfactory (piriform) cortex. In contrast to congenital sensory loss in other systems, we show that morphological alterations due to lifelong olfactory deprivation are limited. Alterations are primarily localized around the olfactory sulcus and likely due to the absence of olfactory bulbs. A possible explanation for the lack of major morphological alterations in individuals with congenital anosmia is that the olfactory regions may be recruited for non-olfactory functions.

---

Sensory deprivation, an extended and significant reduction in (or complete absence of) sensory input, has commonly been associated with altered brain morphology in areas that typically process information from the absent sense (Noppeney et al. 2005; Peter et al. 2020; Simon et al. 2020). These morphological changes are especially pronounced in cases of life-long deprivation where gray matter changes are present in several processing areas (Manno et al. 2021; Grégoire et al. 2022; Lin et al. 2022). Adults who were born without a sense of smell with no other underlying medical conditions (isolated congenital anosmia; ICA) display morphological deviations in the orbitofrontal cortex (Peter et al. 2020; Peter et al. 2023) which has been described as the primary area of advanced olfactory processing (Gottfried and Zald 2005; Lundström et al. 2011). Interestingly, these changes occur mainly outside of the primary olfactory cortex (Peter et al. 2020), with gray matter volume atrophy in the bilateral olfactory sulci and increased gray matter volume in the medial orbital gyri; and no apparent changes in the cortical structure most associated with olfactory processing, the piriform cortex. Previous studies have focused mainly on gray matter morphology (the processing dimension) and significantly less work has been devoted to assessing white matter morphology (the communication dimension); we therefore know substantially less about how olfactory deprivation affects the deeper tissues of the brain.

The effects of sensory loss on white matter have been established in vision and audition, with similar findings of changes in volume in both modalities (Pan et al. 2007; Manno et al. 2021). For instance, congenitally blind individuals express overall a reduction of white matter volume compared to healthy controls, with specific alterations in areas necessary for transfer of visual information between the two hemispheres (Ptito et al. 2008). White matter changes are also prominent in individuals with acquired vision loss, who show more extensive alterations compared to congenitally blind individuals (Wang et al. 2013). Similarly, deaf individuals exhibit alterations in white matter structural parameters, such as reductions in volume (Simon et al. 2020), density (Olulade et al. 2014), and fractional anisotropy (Hribar et al. 2014). Not surprisingly, these alterations occur across both the primary and secondary auditory cortices, mainly in the bilateral superior temporal gyrus (Grégoire et al. 2022). In essence, various studies on visual and auditory deprivation have established an effect on white matter morphology with possible variation depending on the extent and origin of the loss. Yet, the putative effects of olfactory deprivation on white matter morphology remain largely unexplored.

There have been reports of decreased white matter volume in secondary olfactory areas in individuals with anosmia (Peng et al. 2013), particularly in cases of acquired olfactory loss (Manan et al. 2022) resulting from conditions such as COVID-19 (Yildirim et al. 2022) or trauma (Gao et al. 2022). Furthermore, a positive correlation has been observed between the duration of anosmia and the extent of white matter atrophy (Peng et al. 2013). However, a few studies have reported larger volume (Manan et al. 2022), increased density (Frasnelli et al. 2013; Manan et al. 2022), and heightened fractional anisotropy of white mater among individuals with ICA. Previous studies on brain morphology in anosmia, particularly congenital anosmia, report disparate effects in primary olfactory areas such as the piriform cortex. These inconsistencies may be attributed to variability in the duration and etiology of anosmia, as such diversity, both within and across studies, makes it difficult to draw definitive conclusions about the condition. In addition, many studies show weak effects that might be explained by their small sample sizes.

Here, we assessed alterations in white matter morphology in a comparably large cohort of individuals who were born without the sense of smell, compared to a group of healthy controls. In this very same sample, we previously found increased gray matter volume in the bilateral medial orbital gyrus and decreased gray matter volume in the bilateral olfactory sulcus compared to healthy controls (Peter et al. 2020; Peter et al. 2023), and we therefore hypothesized that ICA individuals would demonstrate white matter alterations in areas linking these regions. Moreover, we previously found no significant difference in gray matter volume in the piriform cortex of individuals with ICA compared to healthy controls. This might seem surprising considering the lifelong absence of olfactory input but may be partially explained by increases in visual and auditory processing within the region. Based on this, we expected to find altered white matter volume adjacent to the piriform cortex in individuals with ICA as compared to normosmic controls.

## MATERIALS AND METHODS

### Participants

A total of 98 participants were included in the study, 49 individuals with ICA and 49 controls, matched in terms of sex and age (Table 1). For the ICA group, the inclusion criterion was a self-reported lifelong lack of smell perception without a known underlying condition. Lack of smell perception was confirmed by the olfactory assessment described below. For the control group, the inclusion criterion was a normal sense of smell confirmed by the same olfactory assessment. Out of the 49 individuals in the ICA group, 37 lacked olfactory bulbs bilaterally. The presence of olfactory bulbs in the remaining 12 was either non-determinable or identifiable but visibly hypoplastic (assessed from structural images). Data was collected in two countries, Sweden and the Netherlands. All participants provided written informed consent prior to participation and the study was approved by ethical review boards in both Sweden (2015/1585-31) and the Netherlands (NL69840.081.19).

**Table 1.** Descriptive statistics of participants per group.

|  | ICA (*n* = 49) | Control (*n* = 49) |
| --- | --- | --- |
| **Sex** |  |  |
| Male | 20 | 20 |
| Female | 29 | 29 |
| **Age** |  |  |
| Mean (SD) | 36.3 (11.7) | 36.1 (11.5) |
| **TDI** |  |  |
| Mean (SD) | 10.3 (2.4) | 35.5 (3.5) |

### Procedure

#### Olfactory Assessment

To ensure anosmia amongst the ICA group and normal olfactory function in the control group, olfactory function was assessed in all participants using the Sniffin’ Sticks extended test battery (Burghart Messtechnik GmbH, Holm, Germany). The test is a validated and standardized assessment of olfactory ability (Hummel et al. 1997; Kobal et al. 2000; Sorokowska et al. 2015) consisting of three subtests measuring odor detection threshold (T), odor quality discrimination (D), and odor quality identification (I), yielding a combined (TDI) score of objective olfactory function (max score of 48). Using normative data from over 9000 people (Oleszkiewicz et al. 2019), individuals can be classified as either anosmic, hyposmic, or normosmic. In our sample, based on the individual’s TDI score and age, all individuals in the ICA group were classified as anosmic and all individuals in the control group as normosmic.

MRI Brain imaging data were collected using Siemens Magnetom 3T MR scanners (Siemens Healthcare, Erlangen, Germany). More specifically, two Prisma scanners using 20-channel head coils and one Verio scanner using a 32-channel head coil was used. T1-weighted images covering the entire brain were acquired using either a 3D GR/IR T1-weighted sequence (208 slices, TR = 2300 ms, TE = 2.89 ms, FA = 9°, voxel size = 1 mm3, and FoV = 256 mm) or an MP-RAGE sequence (176 slices, TR = 1900 ms, TE = 2.52 ms, FA = 9°, voxel size = 1 mm3, and FoV = 256 mm).

### Analysis

#### Voxel-Based Morphometry

To segment T1-weighted images into gray matter, white matter, and cerebrospinal fluid in native space, voxel-based morphometry (Ashburner and Friston 2000) was conducted using the SPM12 software. Gray and white matter segmented images were fed into the DARTEL toolbox (Ashburner 2007) for alignment of the gray and white matter of all participants and generating a brain template through an iterative alignment process. All segmented images and the generated template were then normalized to MNI space using 12-parameters affine normalization. The resulting images were then modulated by the Jacobian determinants of the deformation fields and smoothed using a 6 mm Gaussian kernel.

Voxel-wise group differences in white matter volume between the ICA and control groups were assessed using an independent-samples t-test with age, sex, and data collection site as nuisance covariates. A masking threshold of .15 was applied to exclude non-white matter voxels. The voxel-wise group comparison was finally assessed at family-wise error (FWE)-corrected p < .05.

#### Searchlight Classification

A searchlight classification approach was applied to investigate whether white matter volume patterns could distinguish individuals with ICA from the controls.

Using the CoSMoMVPA toolbox (Oosterhof et al. 2016), we first segmented the white matter images into spherical clusters with a 6 mm radius. Each cluster was then analyzed using a linear support vector machine (SVM) classifier, trained to classify participants as either ICA or controls. To ensure robust classification, we employed a 10-fold cross-validation strategy, training the classifier on 80% of participants and testing it on the remaining 20%, maintaining a balanced distribution across groups. Classification accuracy for each cluster was then mapped onto the center voxel of the corresponding sphere. This process was repeated across the entire brain, resulting in a whole brain classification accuracy map. Clusters with classification accuracy greater than 80% were subsequently evaluated for statistical significance. To accomplish this, 5000 permutations were performed and observed accuracy exceeding 95% of the accuracy of permuted data was considered significant (*p* < .05).

## RESULTS

### Orbitofrontal White Matter Reduction in Isolated Congenital Anosmia

First, we assessed ICA-related alterations in white matter volume with a voxel-wise analysis on segmented white matter maps. We found significantly reduced white matter volume in the bilateral orbitofrontal region in ICA individuals compared to control individuals (*P* _FWE-corrected_ < .009; Figure 1, Table 2). Additionally, we used a more liberal threshold (*P* _uncorrected_ < .001) to assess potential differences between ICA and control individuals in the piriform cortex but found no significant effect extending to this region.

**Table 2.**
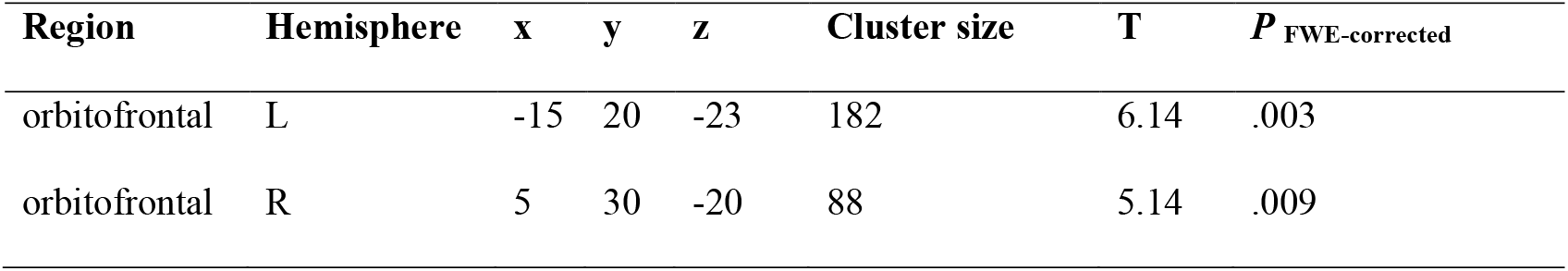
Significant areas of difference between ICA and control individuals. *P* _FWE-corrected_ denotes Family Wise Error corrected p-values.

**Figure 1.**
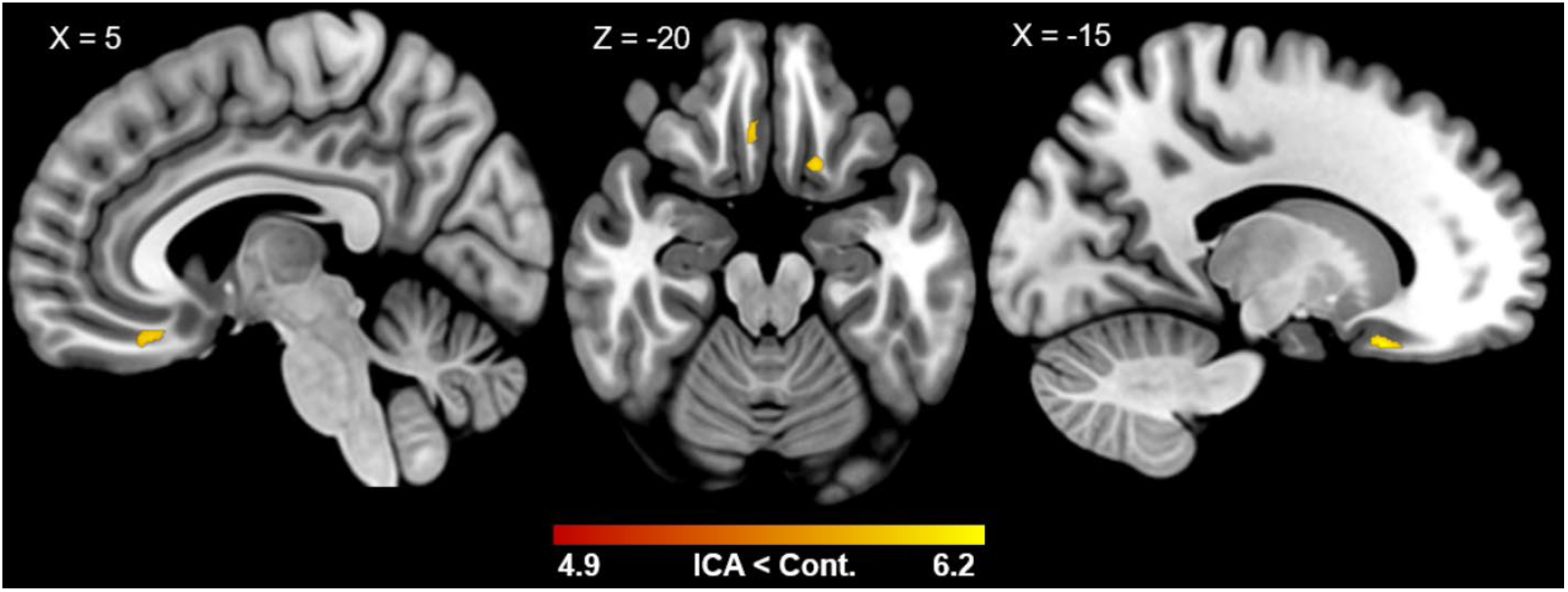
White matter reduction in isolated congenital anosmia (ICA). Individuals with ICA showed reduced white matter volume bilaterally in the orbitofrontal region compared to controls (*P* _FWE-corrected_ < .009). Colors indicate *t*-values.

### Searchlight Classifier Distinguished between ICA group and Controls

Given that voxel-based assessments indicate differences in individual voxels, we wanted to assess also whether summated changes across neighboring structures would be more sensitive to potential changes. Based on the observed altered white matter volume in ICA individuals, we hypothesized that summated white matter volume could distinguish ICA individuals from controls. To test this, we applied a searchlight SVM classifier. The classifier identified two clusters (Figure 2, Table 3) with an accuracy threshold > .80, centered around the bilateral orbitofrontal region that significantly distinguished ICA individuals from normosmic controls. A permutation test confirmed the statistical significance of these clusters (*p* < .0002).

**Table 3.** Areas significantly distinguished ICA individuals from controls.

| Region | Hemisphere | Cluster size | peak coordinates (x, y, z) | peak accuracy |
| --- | --- | --- | --- | --- |
| orbitofrontal | L | 2437 | -6, 42, -24 | .90 |
| orbitofrontal | R | 1581 | 8, 20, -9 | .88 |

**Figure 2.**
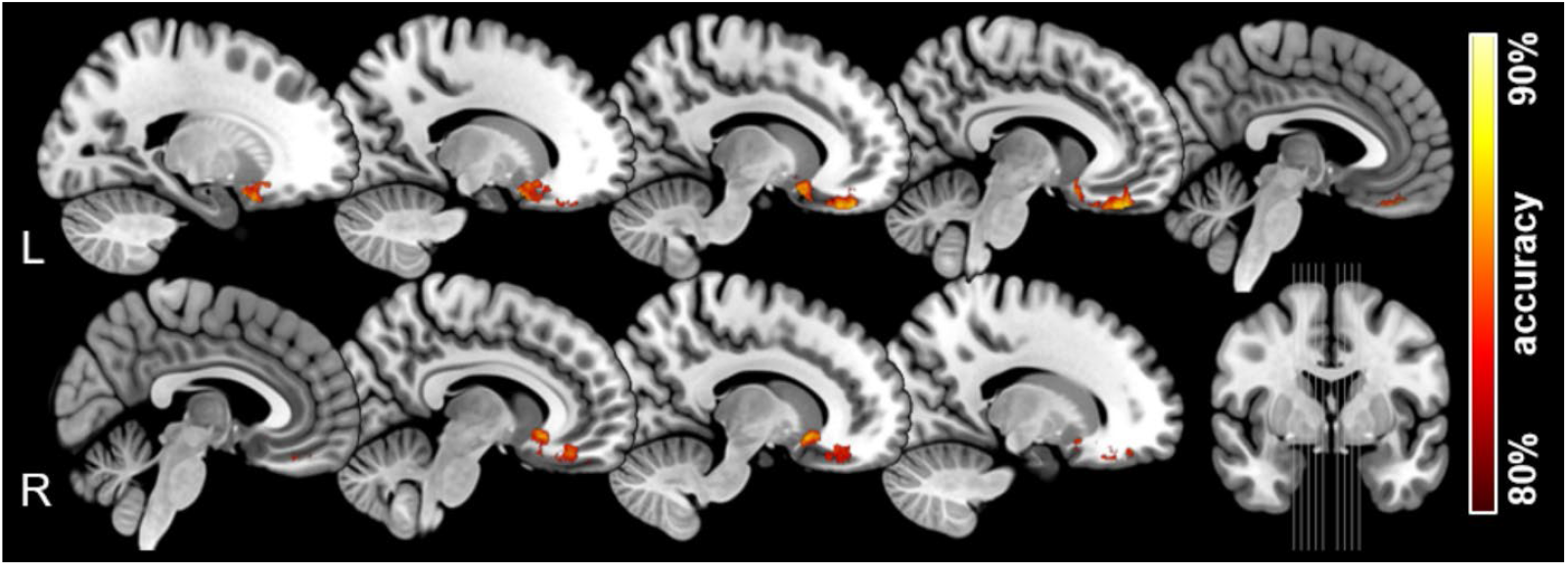
Group classification accuracies between individuals with congenital anosmia and controls, based on white matter volume measures. Significant clusters with accuracy > .80 are located around orbitofrontal region (*p* < .0002). Colors denote percent correct classification between the two groups (accuracy).

## DISCUSSION

We here demonstrate that individuals with isolated congenital anosmia (ICA) have distinct but limited white matter alterations as compared to normosmic controls. Specifically, ICA individuals demonstrate reduced white matter volume bilaterally, in two clusters of the orbitofrontal cortex surrounding the olfactory sulcus. Moreover, we found no significant alterations in the piriform cortex, the primary sensory cortex for olfaction, even when using a liberal threshold. These results are consistent with our previous findings of alterations in gray matter (Peter et al. 2020; Peter et al. 2023), where morphological changes are restricted to the orbitofrontal cortex, indicating that congenital olfactory sensory deprivation yields limited structural reorganization.

Contrary to the large-scale white matter changes observed in individuals with lifelong visual and auditory sensory loss (Pan et al. 2007; Manno et al. 2021), our findings suggest that despite lifelong olfactory deprivation, cerebral morphological alterations are limited. In other words, there seems to be no extensive neural reorganization following lifelong absence of sensory input in the olfactory system. The changes that we do observe around the olfactory sulcus are likely due to the absence of olfactory bulbs in individuals with ICA. The lack of effect in the primary sensory region, taken together with previous findings on gray matter, suggests that sensory deprivation does not affect the olfactory system in the same substantial way as for our other sensory systems (Peter et al. 2023). Potentially, the lack of obvious effects could be linked to the fact that the olfactory system differs from other sensory systems in that the primary sensory cortex (the piriform cortex) is involved in more complex processing and consists of a different type of cortical tissue compared to the other primary sensory cortices (Gottfried 2010). In addition, we have recently shown that the piriform cortex processes also non-olfactory visual and auditory stimuli (Thunell et al. 2025 Apr). If the olfactory system is regularly engaged in processing non-olfactory input, particularly in individuals who have never experienced olfactory stimuli, the synaptic pruning mechanism may be absent in the olfactory system of individuals with ICA.

The lack of wide-range morphological changes also raises questions regarding compensatory mechanisms and whether these exist within the olfactory network or not. Further investigations could help clarify whether individuals with ICA exhibit neural adaptations that compensate for their lack of olfactory input. As outlined above, the fact that the identified morphological changes are limited could potentially be attributed to the olfactory cerebral system being occupied by other functions, such as visual or auditory processing, in individuals suffering from olfactory sensory loss. In line with this argument is our recent demonstration (Thunell et al. 2025 Apr) of the piriform cortex processing visual and auditory sensory stimuli, where we also show that in the absence of olfactory-related information or association, ICA individuals display crossmodal transfer of function to the piriform cortex. In other words, the piriform cortex in ICA individuals processes visual and auditory sensory stimuli and is also more attuned to non-olfactory sensory processing than control individuals with a normal sense of smell (Thunell et al. 2025 Apr). The increased sensory processing in piriform cortex of visual and auditory stimuli could, potentially, compensate for the loss of olfactory-related processing that would have resulted in reduced white matter. Alternatively, the olfactory system might be more robust than our other sensory systems against morphological alterations. Speaking against this alternative explanatory model is the fact that the healthy human olfactory system shows large neuroplasticity where, for example, olfactory bulb morphology is altered by both changes in the amount of odor sensations (Huart et al. 2019) and disease states (Thunell et al. 2022). The olfactory system is also one of the few places where adult neurogenesis has been demonstrated (Nissant et al. 2009). It is therefore likely that the olfactory system’s involvement in amodal sensory processing is the most convincing explanation of the lack of clear and widespread effects. Further research is warranted to explore the functional implications of these findings and to determine whether olfactory-related brain regions are repurposed for other cognitive or sensory functions in individuals with olfactory sensory deprivation.

Differences in morphological changes related to onset of sensory deprivation have been established in both vision and audition with morphological changes varying depending on whether sensory loss is congenital or acquired. With the current sample, we can only establish effects of a lifelong absence of olfactory input and cannot definitively conclude that the same results would emerge in individuals with acquired smell loss, caused by for example viral infections or trauma.

In conclusion, our study demonstrates that congenital anosmia is associated with subtle but specific white matter alterations, primarily localized around the olfactory sulcus. The absence of widespread white matter changes suggests that the olfactory system exhibits more limited structural reorganization in response to congenital sensory deprivation than in the visual and auditory systems.

## AUTHORSHIP CONTRIBUTION

JNL contributed to the conceptualization and design of the study. MP and ET collected data. FD performed the formal analyses. ALW wrote the initial draft of the manuscript. All authors contributed to manuscript revision, as well as read and approved the submitted version.

